# Local fat content determines global and local stiffness in livers with simple steatosis

**DOI:** 10.1101/2022.07.14.500092

**Authors:** David Li, Paul A. Janmey, Rebecca G. Wells

**Author notes:** Correspondence: Rebecca G. Wells, 905 BRB II/III, 421 Curie Boulevard, Philadelphia, PA 19104-6140. Data Transparency: All study data are included in the article and/or supplementary material, and all analytic methods are available upon request from the authors.

## Abstract

Fat accumulation during liver steatosis precedes inflammation and fibrosis in fatty liver diseases, and is associated with disease progression. Despite a large body of evidence that liver mechanics play a major role in liver disease progression, the effect of fat accumulation by itself on liver mechanics remains unclear. Thus, we conducted ex vivo studies of liver mechanics in rodent models of simple steatosis to isolate and examine the mechanical effects of intrahepatic fat accumulation, and found that fat accumulation softens the liver. Using a novel adaptation of microindentation to permit association of local mechanics with microarchitectural features, we found evidence that the softening of fatty liver results from local softening of fatty regions rather than uniform softening of the liver. These results suggest that fat accumulation itself exerts a softening effect on liver tissue. This, along with the localized heterogeneity of softening within the liver, has implications in what mechanical mechanisms are involved in the progression of liver steatosis to more severe pathologies and disease. Finally, the ability to examine and associate local mechanics with microarchitectural features is potentially applicable to the study of the role of heterogeneous mechanical microenvironments in both other liver pathologies and other organ systems.

## Introduction

Liver steatosis is the accumulation of fat within the liver and is associated with progression to more severe pathologies including steatohepatitis, fibrosis, cirrhosis, and hepatocellular carcinoma (1, 2). Affecting over 32% of adults in the US (3), steatosis is among the first pathological changes in both alcoholic and non-alcoholic fatty liver disease, preceding other pathologies such as inflammation and fibrosis that stiffen the liver and often preceding symptoms (1, 4, 5). The effect of fat accumulation itself, in the absence of fibrosis or inflammation, on disease progression, is, however, relatively poorly understood.

In particular, despite the well documented connection between liver stiffness and fibrosis (6-8), the effect of isolated steatosis on the mechanical properties of the liver has yet to be defined. Previous investigations in other tissues have shown that fat accumulation in tissues alters tissue mechanics, but whether it exerts a softening or stiffening effect has been variable and tissue specific (9-12). Most studies of liver mechanics during fatty liver disease have focused on later stages of the disease, during which inflammation and fibrotic remodeling may obscure the contribution of fat accumulation (12-14). Furthermore, past work was largely carried out in vivo (15-19), where changes in fluid pressure presented a confounding factor (20-22). Thus, the impact of fat accumulation in isolation on liver mechanics is not known.

In addition, fatty liver disease is histologically heterogeneous at the sub-millimeter (mesoscale) (4, 23), which is the scale most relevant to mechanosensing (24-26). This local variation in fat accumulation may cause changes in the mechanical microenvironment of liver cells in ways distinct from that of whole organ mechanics. Due to limitations in methodology, previous work has been unable to link mesoscale architecture and mechanics in the fatty liver. The resolution of ultrasound-based methods such as transient elastography is poor at sub-millimeter scales (27, 28), while atomic force microscopy (AFM) is best at the nano-scale (29). Microindentation provides accurate mechanical measurements at the relevant scale, but it has been difficult to correlate mechanics and architecture (29).

As a result, although there is histological evidence of local heterogeneity in fat accumulation in liver, how this heterogeneity affects the mechanical microenvironment at cell-relevant scales has not been explored.

In this work, we address these questions by determining the effects of simple steatosis, without inflammation or fibrosis, on the mechanics of both the whole liver and heterogeneous regions of fat accumulation within the tissue. Using the ob/ob mouse as a genetic model of simple steatosis, we measured whole liver mechanics by rheometry. The liver was studied ex vivo to avoid confounding from fluid pressure. To determine whether local differences in fat accumulation cause local changes in tissue mechanics, we used a new method to measure local stiffness values by microindentation and to demarcate the measured regions for subsequent correlation with lipid droplet staining and other microarchitectural studies. With this new combined approach, we determined that fat accumulation softens the liver at both whole organ and meso scales. This may have important implications for understanding the progression of fatty liver disease.

## Materials and Methods

### Animal studies

All animal work was carried out in strict accordance with the recommendations in the Guide for the Care and Use of Laboratory Animals of the National Institutes of Health. Animal protocols were approved by the Institutional Animal Care and Use Committee of the University of Pennsylvania (protocol #804031). Ob/ob mice were obtained from the Jackson Laboratories (strain #000632) and were housed in a temperature-controlled environment with appropriate enrichment, ad libitum feeding of standard rodent chow and water, and 12h light/dark cycles. Euthanasia was carried out by CO_2_ inhalation followed by exsanguination. Livers were harvested from ob/ob mice and wild type littermates at 8w, 12w, and 36w and stored in PBS at 4°C for up to 5h before being analyzed; rodent livers are adequately preserved under these conditions and rheological properties are maintained (30, 31).

### Shear rheometry

Shear rheometry was performed as described previously (30). Briefly, liver samples were prepared using an 8 mm punch (MP0144, Alabama R&D) with all punches taken in the same orientation from the same lobe. The height of the slices ranged from 2.0 to 3.9 mm in the uncompressed state. Samples were kept hydrated during all experiments with PBS. Parallel plate shear rheometry was carried out on a Kinexus PRO rheometer (Kinexus series, Malvern Instruments) at room temperature. Samples were attached to rheometer platforms with fibrin glue by applying 5 μL each of 20 mg/mL bovine plasma fibrinogen (341573, Sigma-Aldrich) and 100 U/mL bovine thrombin (T4648, Sigma-Aldrich) to both the top and bottom sides of the sample. The upper platen was quickly lowered until 0.02 N of nominal initial force was applied to ensure adhesive contact of the sample with the metal surfaces of the rheometer, and the sample was allowed to sit for 10 min to allow the fibrin glue to polymerize fully before performing measurements. The fibrin glue does not affect the mechanics of the system (30). Mechanics were measured with a dynamic time sweep test (2% constant strain, oscillation frequency 1rad/s, measurements taken for 120s). These measurements were carried out first uncompressed, then with increasing uniaxial tension (10% and 20%), then uncompressed, then with increasing uniaxial compression (10, 15, 20, and 25%). Tension and compression were applied by changing the gap between the platform and upper platen of the rheometer. The following correction was applied to account for the change in cross-sectional area during testing under the assumption that total tissue volume is conserved, where G’ is the storage modulus and λ is axial strain:

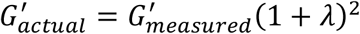

A similar correction was applied to the loss modulus G’’ and normal stress δ. Young’s modulus E was determined by calculating the slope of stress (Pa) vs axial strain (30). Young’s modulus E at zero compression was determined by the slope of stress vs axial strain between -10% and 10% compression.

### Microindentation and demarcation of microindented regions

Microindentation was performed as described previously (29), with additional modifications to permit the demarcation and visualization of mesoscale features within the microindented regions of the tissue (Figure 1). Briefly, the microindentation device consists of a stepping motor (L4018S1204-M6, Nanotec) attached to a μN-resolution tensiometric probe adapted from the surface tension measurement apparatus of a Langmuir monolayer trough (MicroTrough X, Kibron Inc), consisting of a 0.510 mm diameter blunt-ended cylindrical tungsten alloy wire hung from a digital microbalance. The force-displacement relationship was found to behave as a Hookean spring with force linearly related to displacement, and the spring constant was calibrated before each (k_probe_ ∼ 4.5N/m).

**Figure 1.**
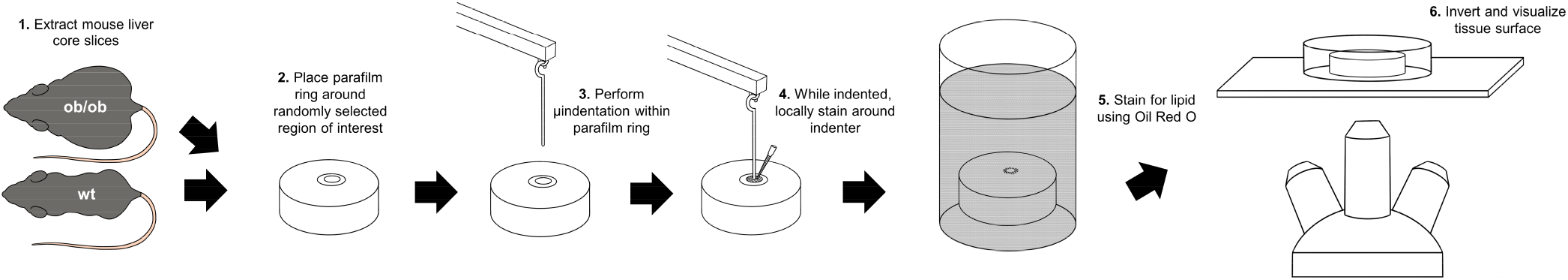
Schematic of technique combining microindentation with visualization of tissue features. After removing livers from mice and generating liver core slices (step 1), parafilm rings were placed around randomly-selected regions of interest on each slice (step 2) and microindentation was performed on the region within each ring (step 3). After measuring local mechanics, the indenter was lowered to fully indent the microindented region and a stain was locally applied within the parafilm ring around the microindenter (step 4). Afterwards, the tissue slices were stained for lipid using Oil Red O (step 5) and inverted on an epifluorescent microscope to visualize lipid accumulation within each locally-stained region (step 6).

Liver samples 3 mm thick were prepared from liver tissue cores obtained using an 8 mm punch, removing the liver capsule. A ring of Parafilm was prepared using a 2 mm disposable biopsy punch and placed around a randomly selected region of interest on top of the liver sample. Samples were then manually positioned under the free-hanging probe, followed by downward displacement of the probe until contact occurred between the probe and the sample in the center of the Parafilm ring. Following establishment of contact between probe and sample, the probe was translated downward at a continuous rate of 0.0125 mm/s. These translations resulted in decreases in measurable force from the probe, which were converted into local Young’s modulus as described previously (29). After the probe was fully indented into the sample (as determined by the absence of measurable force from the probe), 0.5 μL of 100 μg/mL DAPI (D1306, Invitrogen) solution was pipetted around the probe and allowed to selectively stain tissue within the Parafilm ring for 15s before the sample was washed with PBS. Multiple measurements were obtained per liver sample using this approach. After microindentation and demarcation, liver samples were stored in PBS at 4°C for up to 5h before being stained and imaged for lipid accumulation.

### Visualization and quantification of lipid accumulation of microindented regions

Liver samples were stained for lipid using an adaptation of whole mount Oil Red O staining (32). All incubation steps were performed at room temperature on a shaker. Briefly, samples were incubated with 0.5% Tween 20 (#170-6531, Bio-Rad) in PBS for 15min, changing the solution every 5min, before incubating with 0.5% Oil Red O in propylene glycol (D1306, Poly Scientific R&D Corp) for 15min.

Afterwards, the samples were washed once with Tween solution before incubating with Tween solution for 15min, changing the solution every 5min. Whole mount Oil Red O staining of liver slices served as a rapid and easy technique to visualize the heterogeneous distribution of lipid on the surface of liver tissue samples (Figure 2A).

**Figure 2:**
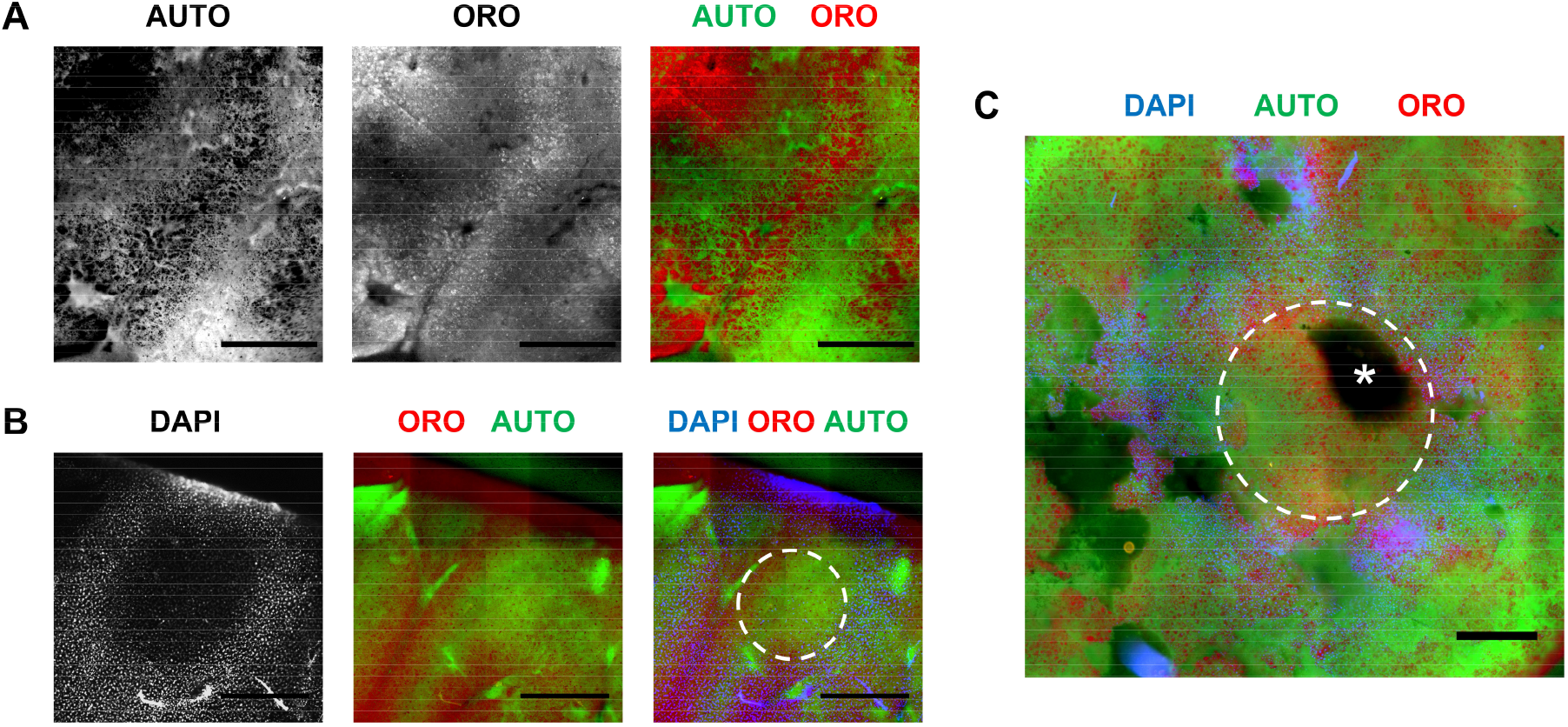
Whole mount fluorescent staining and microscopy shows features of ob/ob livers. (A) Whole mount epifluorescence microscopy images of autofluorescence (left) and intracellular lipid accumulation in hepatocytes show spatial heterogeneity of lipid accumulation at the submillimeter scale of 12w ob/ob livers. (B) Whole mount epifluorescence microscopy images of a 12w wt liver showing DAPI (blue) demarcating the microindented region (left), autofluorescence (green) and heterogeneous lipid accumulation using Oil Red O (red, middle), and the three channels merged (right) showing low lipid in the indented region (white dashed line). (C) Epifluorescent composite image of a 12w ob/ob liver with region of indentation from a 1mm indenter marked with DAPI and stained for lipid accumulation (DAPI, blue; autofluorescence, green; lipid, red). * indicates an intrahepatic blood vessel within the indented region (white dashed line). Scale bars = 500 μm.

Visualization of lipid accumulation in microindented regions was performed using whole mount epifluorescence microscopy on a Nikon Eclipse Ti widefield microscope equipped with a 4x/0.31 N.A. PlanFluor dry objective lens and motorized stage. Lipid (33), DAPI demarcation, and liver autofluorescence were visualized using Cy3, DAPI, and FITC filters, respectively. Using this approach, it was possible to visualize both the microindented regions and local lipid accumulation in the same regions (Figure 2B). Using autofluorescence, it was also possible to identify microscale anatomical features on the surface of the tissue such as intrahepatic blood vessels (Figure 2C). Quantification of lipid accumulation within demarcated DAPI rings was performed by simple thresholding of the Cy3 channel after shading correction in ImageJ.

### Statistical analysis

The statistical significance of differences between strain-dependent shear rheometry curves (G’, G’’, and E) of ob/ob mice and wild type mice was determined using two-way ANOVA (30). Statistical significance of the decreasing monotonic trend between increasing local lipid accumulation and local stiffness within sub-mm regions of the ob/ob and wild type mouse liver was determined using Spearman’s rank correlation coefficient (34). For all statistical analyses, a P value of ≤ 0.05 was considered significant.

## Results

### Simple steatosis is associated with liver softening

To determine the effect of fat accumulation on the solid mechanical properties of the liver without influence from other effects such as variations in perfusion, we compared steatotic and non-steatotic mouse livers ex vivo by parallel plate rheometry. We used ob/ob mice as a model of liver steatosis without inflammation or fibrosis, with wild-type littermates as controls. Livers were removed at 8, 12, and 36 weeks of age; H&E staining showed an increase in lipid droplets in ob/ob mice at all ages compared to controls (Figure 3A). Notably, in ob/ob mice at 36 weeks of age, both the area of fatty liver and the number of visible droplets had decreased compared to ob/ob mice at 12 weeks of age.

**Figure 3.**
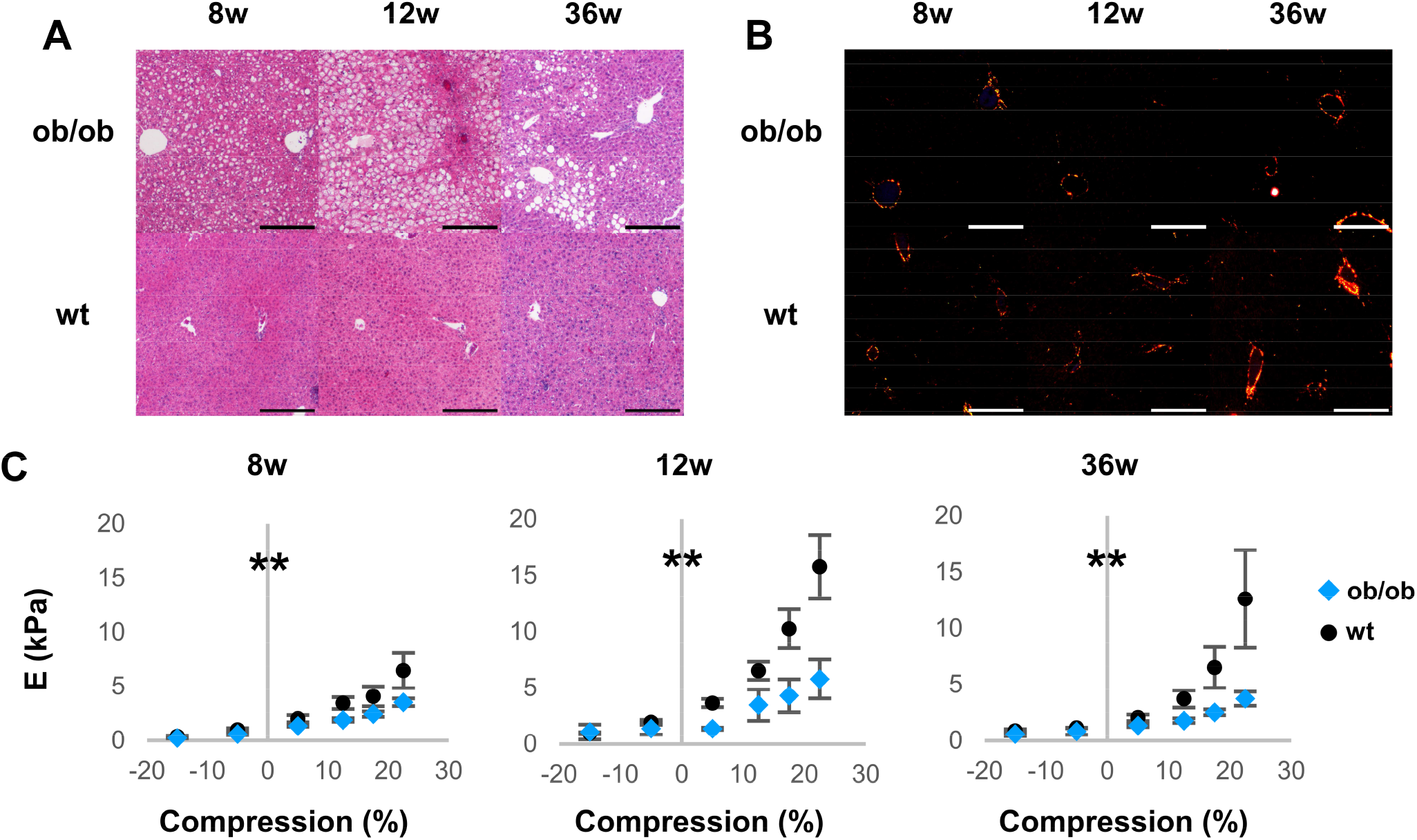
Simple steatosis is associated with lower Young’s modulus in ob/ob mice. (A) Representative H&E stains of livers from ob/ob mice and wild type littermates taken at 8w, 12w, and 36w of age. (B) Representative picrosirius red stains of ob/ob and wt livers taken under polarized light microscopy. Scale bar = 250 μm. (C) Young’s modulus E of ob/ob (blue) and wt (black) livers at 8w, 12w, and 36w under varied tension and compression. Ob/ob livers were significantly softer than wt livers. n ≥ 4 animals for each condition. Error bars indicate standard error. ** P < 0.005 for differences between curves by two-way ANOVA.

There was no evidence of inflammation in either ob/ob or control livers, and picrosirius red staining for collagen confirmed the absence of fibrosis in both (Figure 3B and Supplementary Figure 1).

Shear rheometry was performed on liver cores (7) to determine the Young’s modulus E, showing that livers from ob/ob mice were significantly softer and had less pronounced stiffening under physiologically relevant degrees of compression (30) than those from age-matched controls (Figure 3C). The Young’s moduli E at zero compression was extrapolated from the curve, and was found to be significantly softer in ob/ob mice at 8w and 12w than those from age-matched controls (1.50±0.20 vs. 0.98±0.07 kPa for 8w and 2.78±0.28 vs. 1.36±0.30 kPa for 12w, P < 0.05 for 8w and 12w by Student’s T-test). However, there was no significant difference between E at zero compression at 36w, when the differences in lipid accumulation were reduced between livers of ob/ob mice and age-matched controls (1.59±0.14 vs. 1.09±0.20 kPa for 36w, no significance by Student’s T-test). These results suggest that the degree of steatosis, rather than the duration of steatosis or age, is associated with liver softening.

The liver is subject to shear in addition to compressive forces in vivo (30), and so shear rheometry was also used to determine the shear storage modulus G’ and shear loss modulus G” of ob/ob and wild type liver cores. Although the liver was viscoelastic for all conditions, G” was always significantly lower than G’ and contributed minimally to the complex shear modulus. With the exception of livers at 36w under zero compression or tension, all livers from ob/ob mice had significantly lower moduli and less-pronounced compression stiffening than those from age-matched controls (Figure 4). Thus, ex vivo mechanical characterization of ob/ob and wild type mouse livers demonstrates that steatosis is associated with overall liver softening.

**Figure 4.**
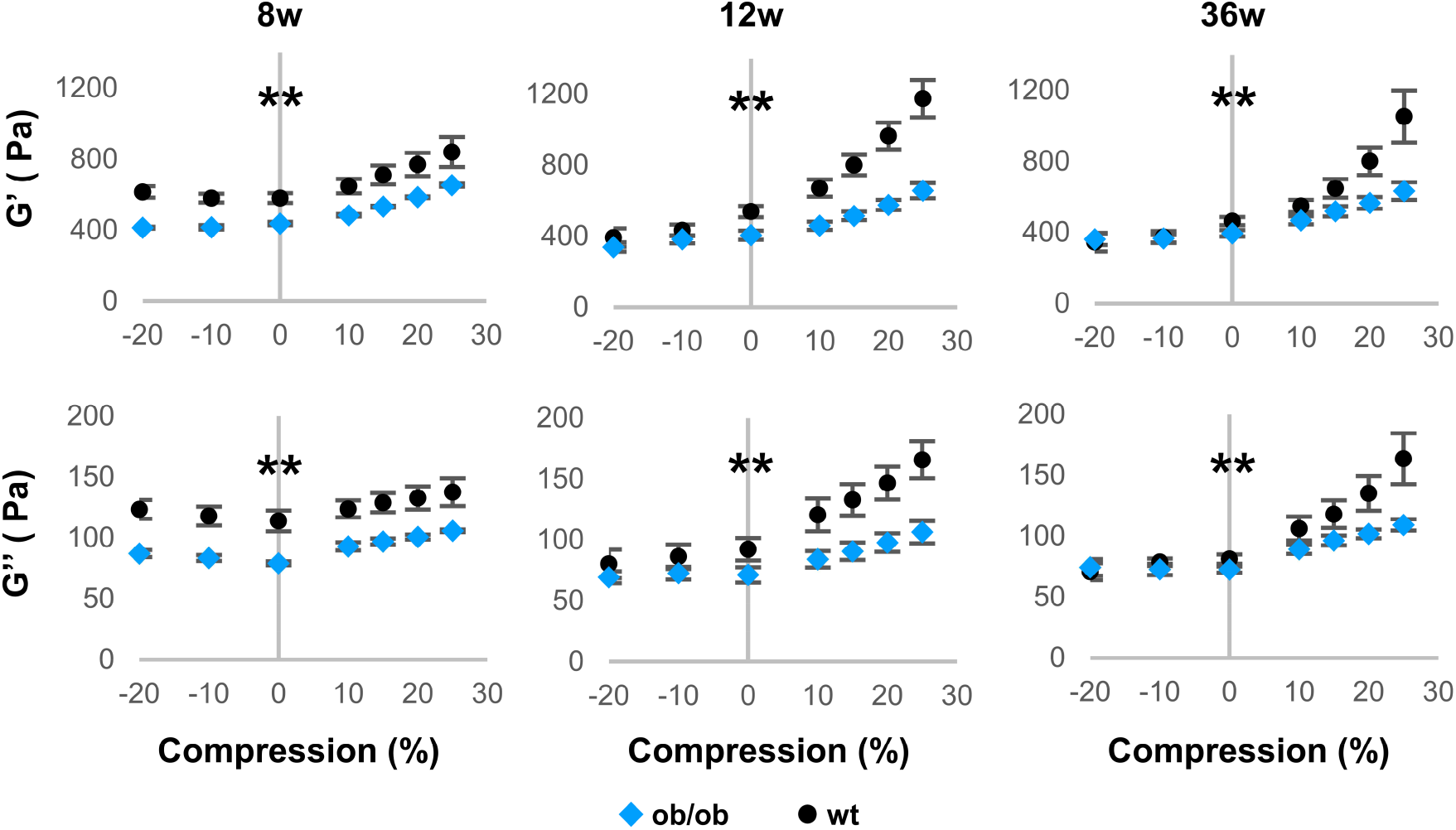
Simple steatosis is associated with lower shear elastic and loss moduli in ob/ob mice. Shear elastic modulus G’ (top) and loss modulus G’’ (bottom) of ob/ob (blue) and wt (black) livers at 8w, 12w, and 36w under varied tension and compression. Ob/ob livers had significantly lower shear elastic and loss moduli than wt livers and less compression stiffening. n ≥ 4 animals for each condition. Error bars indicate standard error. ** P < 0.005 for differences between curves by two-way ANOVA.

### Association of microindentation with visualization of histological features

Shear rheometry measures the overall mechanics of tissue cores which are millimeters in diameter, and thus is limited to measuring stiffness on the same spatial scale as conventional clinical ultrasound and MRE approaches (27, 28). These measurements do not capture variations in stiffness due to features at the micron scale. For example, fat accumulation is heterogeneous in liver steatosis, preferentially localizing to regions around the central veins (23). We therefore developed a technique to determine whether localized lipid accumulation is associated with localized softening.

We adapted a previously described ex vivo microindentation technique (29) to add the capability of associating cell and tissue structures such as lipid droplets to stiffness measurements within the same submillimeter regions of interest (Figure 1). DAPI was applied to liver cores after maximal indentation, staining around but not under the indenter, resulting in a ring demarcating the region where stiffness was measured. After release of the indenter, whole mount Oil Red O staining was used to label lipids. We found that epifluorescence microscopy could be used to simultaneously capture the ring of demarcation by DAPI, lipid accumulation, and liver autofluorescence (Figure 2A and B). Liver autofluorescence was used to avoid measurements over regions with large blood vessels. (Figure 2C).

### Localized fat accumulation is associated with local softening

We used this adapted microindentation technique to determine the relationship between local lipid accumulation and local stiffness measured ex vivo in ob/ob and wild type mouse livers in the absence of other influences such as fluid perfusion. The amount of fat within the demarcated regions was greater in ob/ob livers than controls, and there was marked variation between different demarcated regions in ob/ob livers (Figure 5A). We quantified the amount of lipid staining in the demarcated microindented regions and found a significant inverse correlation between fat content and local stiffness through Spearman’s rank correlation across all ages (Figure 5B). The 12w samples, which had the highest lipid accumulation, demonstrated a plateau (percolation threshold) at approximately 50-60% lipid, beyond which no further softening was observed. Thus, local regions of simple steatosis in rodent livers resulted in associated local softening.

**Figure 5.**
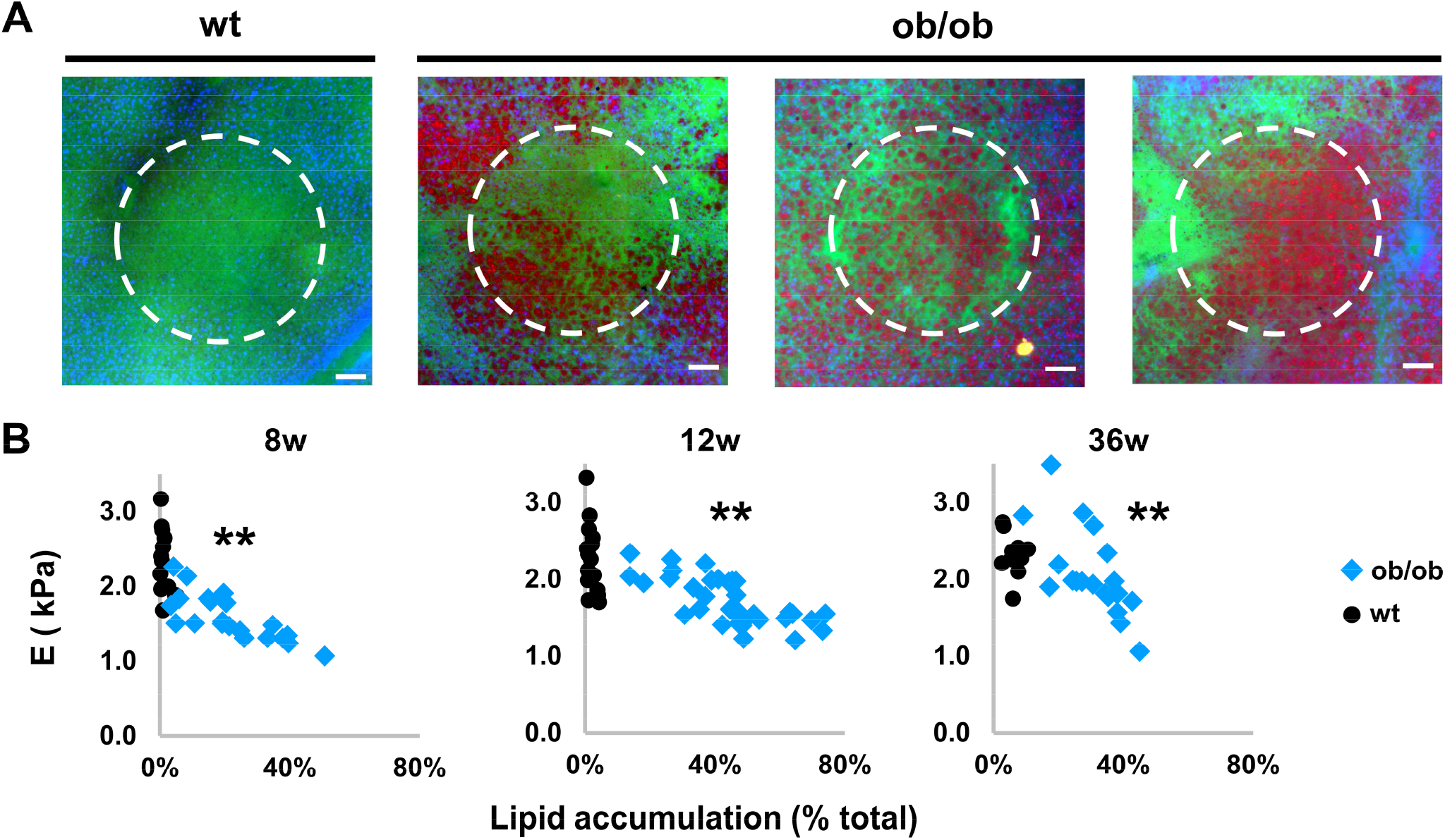
Increased local lipid accumulation in steatosis is associated with local softening. (A) Representative fluorescent images of indented wt and ob/ob livers with increasing lipid accumulation within the indented region, showing DAPI (blue) demarcating the microindented regions (white dashed line), autofluorescence (green) and heterogeneous lipid accumulation using Oil Red O (red). Scale bar 100 μm. (B) Scatter plots of Young’s modulus E vs. lipid accumulation within microindented regions of ob/ob (blue) and wt (black) livers taken at 8w, 12w, and 36w. Submillimeter regions of ob/ob livers with greater local steatosis had lower local stiffness at all tested ages. n ≥ 11 measurements from ≥ 3 animals for each condition. ** rs < -0.7 and P < 0.005 by Spearman’s rank correlation.

## Discussion

In this study, we examined the effect of isolated lipid accumulation on liver tissue solid stiffness in the absence of complicating influences in fatty livers such as fibrosis, inflammation, or variations in blood flow (14, 20-22). Using ex vivo approaches to measure the stiffness of steatotic and normal mouse livers at the millimeter and sub-millimeter length scales, we showed that the liver softens substantially in simple steatosis. Furthermore, we also developed a technique to visualize local histology in regions of microindentation and used it to demonstrate that the local accumulation of lipid droplets within steatotic livers is associated with local softening. Together, these results show that intrahepatic fat causes liver softening and that its uneven distribution in livers with simple steatosis generates a mechanically heterogeneous environment at the sub-millimeter scale.

A major advantage of this study is the ex vivo characterization of liver mechanics without confounding from perfusion. Liver steatosis can cause decreased liver perfusion (22, 35-37) and increased transhepatic pressures (21, 38) – both of which can influence measured stiffness in clinical elastography approaches (20, 39). However, changes in stiffness and fluid pressures present different mechanical stimuli to cells, which then cause cells to mount different responses (40-42). As such, characterizing the stiffness in isolation is important to understanding what stimuli are present in steatosis, and in the development of models to investigate the mechanical role of fat accumulation in fatty liver disease progression.

Steatosis-associated softening at the whole liver level did not automatically mean that steatotic regions would be soft – an alternative hypothesis was that softening occurred preferentially in non-fatty regions of the steatotic liver, possibly due to ECM remodeling (43). Our new demarcated microindentation-microscopy technique was crucial in determining that localized fat accumulation caused local softening in the lipid-laden regions. This finding has important implications in understanding the mechanical contributions of fat accumulation to liver disease given that local transitions in and of themselves can serve as mechanical stimuli and cause changes in cell and tissue behavior (25, 42, 44, 45). The technique also enabled us to determine that there was a maximum threshold of local fat content at which softening plateaued, as predicted by computational models of systems with two phases of differing mechanical properties (46, 47). The demarcated microindentation-microscopy technique will be applicable to the study of other microarchitectural features in fatty liver disease (23) such as local regions of inflammation or ECM remodeling, as well as studies of microarchitectural and mechanical heterogeneity in other tissue systems and pathologies.

The fat-associated liver softening we observed could result from a variety of biophysical changes in fatty liver tissues and cells. Fat accumulation may induce changes in collagen crosslinking (7) or ECM composition (30). Since lipid accumulation causes hepatocytes to swell substantially (48), it is also possible that simple steatosis causes a decrease in the amount of total ECM within a given volume of liver and thus decreases both the stiffness and compression stiffening response. Hepatocyte swelling would also cause any given volume of liver to contain less total cell membrane surface area, which has been hypothesized to contribute to strain stiffening in normal and fibrotic livers by acting as an impediment to fluid movement (30).

There are some limitations to this study. All measurements were obtained on a genetic model of simple steatosis, and additional models of simple steatosis such as those from increased dietary sucrose could be considered to determine whether underlying differences in the models result in differences in cell and tissue response (49). Additionally, the mechanics of the fatty liver in some models might vary as a result of differences in the composition of lipid droplets compared to droplet composition in ob/ob mice (50).

In summary, we show that fat accumulation is associated with both global and local liver softening, a finding that may have significant implications for understanding the progression of early stage fatty liver disease. The ability to correlate submillimeter-scale stiffness measurements with histological features can be used to study the mechanical contributions of microarchitectural changes during the progression of fatty liver disease, and to investigate development and disease in other tissues and organ systems.

## Acknowledgements

The authors thank Jessica Llewellyn, Dongning Chen, Xuechen Shi, and Jeff Byfield for training, technical support, and feedback, the Center for Molecular Studies in Digestive and Liver Diseases (P30DK050306) for histology and microscopy support, Guy Genin for thought-provoking discussions, and Yang Liu for assistance with the illustration.

## Supplementary information

### Supplementary Figures

**Supplementary Figure 1:**
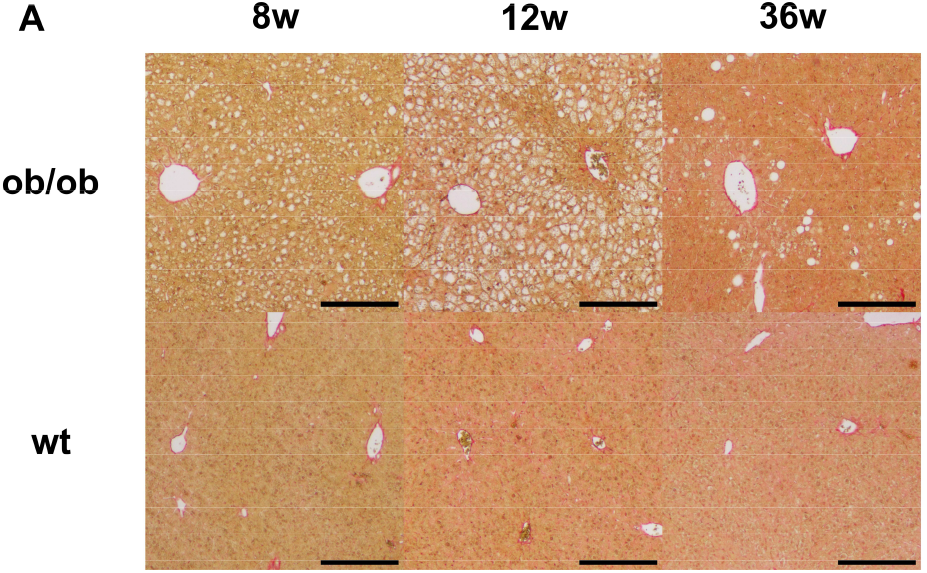
Ob/ob mice do not show fibrosis. Representative picrosirius red stains of ob/ob and wt livers taken at 8w, 12w, and 36w under brightfield microscopy. Images are of the same regions as shown in Figure 3B. Scale bars = 250 μm.

## Abbreviations

NAFLD: non-alcoholic fatty liver disease
NASH: non-alcoholic steatohepatitis
DAPI: 4’,6’-diamidino-2-phenylindole
MRE: magnetic resonance elastography

## Notes

Grant Support: This work was supported by NIH grant R01EB017753 to PAJ and RGW and by the Center for Engineering MechanoBiology (CEMB), an NSF Science and Technology Center, under grant agreement CMMI1548571.

Disclosures: The authors disclose no conflicts.

### Competing Interest Statement

The authors have declared no competing interest.

### Summary of Updates

Expanded figures and discussion.

## References

1. Angulo, P. (2002) Nonalcoholic fatty liver disease. N Engl J Med 346, 1221–1231

2. Chalasani, N., Younossi, Z., Lavine, J. E., Charlton, M., Cusi, K., Rinella, M., Harrison, S. A., Brunt, E. M., and Sanyal, A. J. (2018) The diagnosis and management of nonalcoholic fatty liver disease: Practice guidance from the American Association for the Study of Liver Diseases. Hepatology 67, 328–357

3. Younossi, Z. M., Stepanova, M., Younossi, Y., Golabi, P., Mishra, A., Rafiq, N., and Henry, L. (2020) Epidemiology of chronic liver diseases in the USA in the past three decades. Gut 69, 564–568

4. Theise, N. D. (2013) Histopathology of alcoholic liver disease. Clin Liver Dis (Hoboken) 2, 64–67

5. Matteoni, C. A., Younossi, Z. M., Gramlich, T., Boparai, N., Liu, Y. C., and McCullough, A. J. (1999) Nonalcoholic fatty liver disease: a spectrum of clinical and pathological severity. Gastroenterology 116, 1413–1419

6. Georges, P. C., Hui, J. J., Gombos, Z., McCormick, M. E., Wang, A. Y., Uemura, M., Mick, R., Janmey, P. A., Furth, E. E., and Wells, R. G. (2007) Increased stiffness of the rat liver precedes matrix deposition: implications for fibrosis. Am J Physiol Gastrointest Liver Physiol 293, G1147–1154

7. Perepelyuk, M., Terajima, M., Wang, A. Y., Georges, P. C., Janmey, P. A., Yamauchi, M., and Wells, R. G. (2013) Hepatic stellate cells and portal fibroblasts are the major cellular sources of collagens and lysyl oxidases in normal liver and early after injury. Am J Physiol Gastrointest Liver Physiol 304, G605–614

8. Kostallari, E., Wei, B., Sicard, D., Li, J., Cooper, S. A., Gao, J., Dehankar, M., Li, Y., Cao, S., Yin, M., Tschumperlin, D. J., and Shah, V. H. (2022) Stiffness is associated with hepatic stellate cell heterogeneity during liver fibrosis. Am J Physiol Gastrointest Liver Physiol 322, G234–G246

9. Shoham, N., Girshovitz, P., Katzengold, R., Shaked, N. T., Benayahu, D., and Gefen, A. (2014) Adipocyte stiffness increases with accumulation of lipid droplets. Biophys J 106, 1421–1431

10. Yu, H., Tay, C. Y., Leong, W. S., Tan, S. C., Liao, K., and Tan, L. P. (2010) Mechanical behavior of human mesenchymal stem cells during adipogenic and osteogenic differentiation. Biochem Biophys Res Commun 393, 150–155

11. Labriola, N. R., and Darling, E. M. (2015) Temporal heterogeneity in single-cell gene expression and mechanical properties during adipogenic differentiation. J Biomech 48, 1058–1066

12. Abuhattum, S., Kotzbeck, P., Schlussler, R., Harger, A., Ariza de Schellenberger, A., Kim, K., Escolano, J. C., Muller, T., Braun, J., Wabitsch, M., Tschop, M., Sack, I., Brankatschk, M., Guck, J., Stemmer, K., and Taubenberger, A. V. (2022) Adipose cells and tissues soften with lipid accumulation while in diabetes adipose tissue stiffens. Sci Rep 12, 10325

13. Chin, L., Theise, N. D., Loneker, A. E., Janmey, P. A., and Wells, R. G. (2020) Lipid droplets disrupt mechanosensing in human hepatocytes. Am J Physiol Gastrointest Liver Physiol 319, G11–G22

14. Mueller, S., Millonig, G., Sarovska, L., Friedrich, S., Reimann, F. M., Pritsch, M., Eisele, S., Stickel, F., Longerich, T., Schirmacher, P., and Seitz, H. K. (2010) Increased liver stiffness in alcoholic liver disease: differentiating fibrosis from steatohepatitis. World J Gastroenterol 16, 966–972

15. Wong, V. W., Vergniol, J., Wong, G. L., Foucher, J., Chan, H. L., Le Bail, B., Choi, P. C., Kowo, M., Chan, A. W., Merrouche, W., Sung, J. J., and de Ledinghen, V. (2010) Diagnosis of fibrosis and cirrhosis using liver stiffness measurement in nonalcoholic fatty liver disease. Hepatology 51, 454–462

16. Gaia, S., Carenzi, S., Barilli, A. L., Bugianesi, E., Smedile, A., Brunello, F., Marzano, A., and Rizzetto, M. (2011) Reliability of transient elastography for the detection of fibrosis in non-alcoholic fatty liver disease and chronic viral hepatitis. J Hepatol 54, 64–71

17. Mouzaki, M., Trout, A. T., Arce-Clachar, A. C., Bramlage, K., Kuhnell, P., Dillman, J. R., and Xanthakos, S. (2018) Assessment of Nonalcoholic Fatty Liver Disease Progression in Children Using Magnetic Resonance Imaging. J Pediatr 201, 86–92

18. Petta, S., Maida, M., Macaluso, F. S., Di Marco, V., Camma, C., Cabibi, D., and Craxi, A. (2015) The severity of steatosis influences liver stiffness measurement in patients with nonalcoholic fatty liver disease. Hepatology 62, 1101–1110

19. Hudert, C. A., Tzschatzsch, H., Rudolph, B., Blaker, H., Loddenkemper, C., Muller, H. P., Henning, S., Bufler, P., Hamm, B., Braun, J., Holzhutter, H. G., Wiegand, S., Sack, I., and Guo, J. (2019) Tomoelastography for the Evaluation of Pediatric Nonalcoholic Fatty Liver Disease. Invest Radiol 54, 198–203

20. Millonig, G., Friedrich, S., Adolf, S., Fonouni, H., Golriz, M., Mehrabi, A., Stiefel, P., Poschl, G., Buchler, M. W., Seitz, H. K., and Mueller, S. (2010) Liver stiffness is directly influenced by central venous pressure. J Hepatol 52, 206–210

21. Van der Graaff, D., Kwanten, W. J., Couturier, F. J., Govaerts, J. S., Verlinden, W., Brosius, I., D’Hondt, M., Driessen, A., De Winter, B. Y., De Man, J. G., Michielsen, P. P., and Francque, S.M. (2018) Severe steatosis induces portal hypertension by systemic arterial hyporeactivity and hepatic vasoconstrictor hyperreactivity in rats. Lab Invest 98, 1263–1275

22. Davis, R. P., Surewaard, B. G. J., Turk, M., Carestia, A., Lee, W. Y., Petri, B., Urbanski, S. J., Coffin, C. S., and Jenne, C. N. (2019) Optimization of In vivo Imaging Provides a First Look at Mouse Model of Non-Alcoholic Fatty Liver Disease (NAFLD) Using Intravital Microscopy. Front Immunol 10, 2988

23. Takahashi, Y., and Fukusato, T. (2014) Histopathology of nonalcoholic fatty liver disease/nonalcoholic steatohepatitis. World J Gastroenterol 20, 15539–15548

24. Wang, H., Abhilash, A. S., Chen, C. S., Wells, R. G., and Shenoy, V. B. (2014) Long-range force transmission in fibrous matrices enabled by tension-driven alignment of fibers. Biophys J 107, 2592–2603

25. Wang, Y. L., and Li, D. (2020) Creating Complex Polyacrylamide Hydrogel Structures Using 3D Printing with Applications to Mechanobiology. Macromol Biosci 20, e2000082

26. Stopak, D., and Harris, A. K. (1982) Connective tissue morphogenesis by fibroblast traction. I. Tissue culture observations. Dev Biol 90, 383–398

27. Sandrin, L., Fourquet, B., Hasquenoph, J. M., Yon, S., Fournier, C., Mal, F., Christidis, C., Ziol, M., Poulet, B., Kazemi, F., Beaugrand, M., and Palau, R. (2003) Transient elastography: a new noninvasive method for assessment of hepatic fibrosis. Ultrasound Med Biol 29, 1705–1713

28. Wu, C. H., Ho, M. C., Jeng, Y. M., Hsu, C. Y., Liang, P. C., Hu, R. H., Lai, H. S., and Shih, T. T. (2014) Quantification of hepatic steatosis: a comparison of the accuracy among multiple magnetic resonance techniques. J Gastroenterol Hepatol 29, 807–813

29. Levental, I., Levental, K. R., Klein, E. A., Assoian, R., Miller, R. T., Wells, R. G., and Janmey, P.A. (2010) A simple indentation device for measuring micrometer-scale tissue stiffness. J Phys Condens Matter 22, 194120

30. Perepelyuk, M., Chin, L., Cao, X., van Oosten, A., Shenoy, V. B., Janmey, P. A., and Wells, R. G. (2016) Normal and Fibrotic Rat Livers Demonstrate Shear Strain Softening and Compression Stiffening: A Model for Soft Tissue Mechanics. PLoS One 11, e0146588

31. Tan, K., Cheng, S., Juge, L., and Bilston, L. E. (2013) Characterising soft tissues under large amplitude oscillatory shear and combined loading. J Biomech 46, 1060–1066

32. Kim, S. H., Wu, S. Y., Baek, J. I., Choi, S. Y., Su, Y., Flynn, C. R., Gamse, J. T., Ess, K. C., Hardiman, G., Lipschutz, J. H., Abumrad, N. N., and Rockey, D. C. (2015) A post-developmental genetic screen for zebrafish models of inherited liver disease. PLoS One 10, e0125980

33. Koopman, R., Schaart, G., and Hesselink, M. K. (2001) Optimisation of oil red O staining permits combination with immunofluorescence and automated quantification of lipids. Histochem Cell Biol 116, 63–68

34. Sunderhauf, A., Hicken, M., Schlichting, H., Skibbe, K., Ragab, M., Raschdorf, A., Hirose, M., Schaffler, H., Bokemeyer, A., Bettenworth, D., Savitt, A. G., Perner, S., Ibrahim, S., Peerschke, E. I., Ghebrehiwet, B., Derer, S., and Sina, C. (2021) Loss of Mucosal p32/gC1qR/HABP1 Triggers Energy Deficiency and Impairs Goblet Cell Differentiation in Ulcerative Colitis. Cell Mol Gastroenterol Hepatol 12, 229–250

35. Seifalian, A. M., Chidambaram, V., Rolles, K., and Davidson, B. R. (1998) In vivo demonstration of impaired microcirculation in steatotic human liver grafts. Liver Transpl Surg 4, 71–77

36. Farrell, G. C., Teoh, N. C., and McCuskey, R. S. (2008) Hepatic microcirculation in fatty liver disease. Anat Rec (Hoboken) 291, 684–692

37. McCuskey, R. S., Ito, Y., Robertson, G. R., McCuskey, M. K., Perry, M., and Farrell, G. C. (2004) Hepatic microvascular dysfunction during evolution of dietary steatohepatitis in mice. Hepatology 40, 386–393

38. van der Graaff, D., Chotkoe, S., De Winter, B., De Man, J., Casteleyn, C., Timmermans, J. P., Pintelon, I., Vonghia, L., Kwanten, W. J., and Francque, S. (2022) Vasoconstrictor antagonism improves functional and structural vascular alterations and liver damage in rats with early NAFLD. JHEP Rep 4, 100412

39. Tang, A., Cloutier, G., Szeverenyi, N. M., and Sirlin, C. B. (2015) Ultrasound Elastography and MR Elastography for Assessing Liver Fibrosis: Part 2, Diagnostic Performance, Confounders, and Future Directions. AJR Am J Roentgenol 205, 33–40

40. Olsen, A. L., Bloomer, S. A., Chan, E. P., Gaca, M. D., Georges, P. C., Sackey, B., Uemura, M., Janmey, P. A., and Wells, R. G. (2011) Hepatic stellate cells require a stiff environment for myofibroblastic differentiation. Am J Physiol Gastrointest Liver Physiol 301, G110–118

41. Qi, F., Hu, J. F., Liu, B. H., Wu, C. Q., Yu, H. Y., Yao, D. K., and Zhu, L. (2015) MiR-9a-5p regulates proliferation and migration of hepatic stellate cells under pressure through inhibition of Sirt1. World J Gastroenterol 21, 9900–9915

42. Moriyama, K., and Kidoaki, S. (2019) Cellular Durotaxis Revisited: Initial-Position-Dependent Determination of the Threshold Stiffness Gradient to Induce Durotaxis. Langmuir 35, 7478–7486

43. Kisseleva, T., and Brenner, D. (2021) Molecular and cellular mechanisms of liver fibrosis and its regression. Nat Rev Gastroenterol Hepatol 18, 151–166

44. Lo, C. M., Wang, H. B., Dembo, M., and Wang, Y. L. (2000) Cell movement is guided by the rigidity of the substrate. Biophys J 79, 144–152

45. Lachowski, D., Cortes, E., Robinson, B., Rice, A., Rombouts, K., and Del Rio Hernandez, A. E. (2018) FAK controls the mechanical activation of YAP, a transcriptional regulator required for durotaxis. FASEB J 32, 1099–1107

46. Chen, Y., and Schuh, C. A. (2016) Elasticity of Random Multiphase Materials: Percolation of the Stiffness Tensor. Journal of Statistical Physics 162, 232–241

47. Kantor, Y., and Webman, I. (1984) Elastic Properties of Random Percolating Systems. Physical Review Letters 52, 1891–1894

48. Hall, A., Covelli, C., Manuguerra, R., Luong, T. V., Buzzetti, E., Tsochatzis, E., Pinzani, M., and Dhillon, A. P. (2017) Transaminase abnormalities and adaptations of the liver lobule manifest at specific cut-offs of steatosis. Sci Rep 7, 40977

49. Song, Z., Deaciuc, I., Zhou, Z., Song, M., Chen, T., Hill, D., and McClain, C. J. (2007) Involvement of AMP-activated protein kinase in beneficial effects of betaine on high-sucrose diet-induced hepatic steatosis. Am J Physiol Gastrointest Liver Physiol 293, G894–902

50. Thiam, A. R., Farese, R. V., Jr., and Walther, T. C. (2013) The biophysics and cell biology of lipid droplets. Nat Rev Mol Cell Biol 14, 775–786

